# Modulation of locus coeruleus neurons and strong release of noradrenaline during acute hippocampal seizures in rats

**DOI:** 10.1101/2023.03.06.531292

**Authors:** Lars Emil Larsen, Sielke Caestecker, Latoya Stevens, Pieter van Mierlo, Evelien Carrette, Paul Boon, Kristl Vonck, Robrecht Raedt

**Affiliations:** 4BRAIN, Department of Head and Skin, Ghent University, Ghent, Belgium; Medical Image and Signal Processing, Department of Electronics and Information Systems, Ghent University, Ghent, Belgium; Vrije Universiteit Brussel (VUB), Universitair Ziekenhuis Brussel (UZ Brussel), Department of Medical Oncology/Laboratory for Molecular and Medical Oncology (LMMO), Brussels Belgium.

## Abstract

The locus coeruleus (LC), a brainstem nucleus, is the sole source of noradrenaline in the neocortex, hippocampus and cerebellum. Noradrenaline is a powerful neuromodulator involved in the regulation of excitability and plasticity of large-scale brain networks. In this study, we assessed the activity of locus coeruleus neurons and changes in noradrenergic transmission during acute hippocampal seizures evoked with perforant path stimulation. LC neurons were recorded in anesthetized rats using a multichannel electrophysiology probe and were identified based on electrophysiological characteristics or optogenetic tagging. The majority of LC neurons (55%) were inhibited during seizures, while only a subset of LC neurons (28%) was excited during seizures. Topographic analysis of multi-unit activity showed anatomical separation of neurons that were excited and inhibited during seizures. Changes in hippocampal noradrenaline transmission during seizures were assessed using a fluorescent biosensor for noradrenaline, GRAB_NE2m_, in combination with fiber photometry in both anesthetized and awake rats. Our results indicate that acute electrically evoked hippocampal seizures are associated with strong changes in LC unit activity and strong and consistent time-locked release of noradrenaline. Understanding the role of mass release of noradrenaline during hippocampal seizures is likely to be important to understand seizure pathophysiology.

## Introduction

The locus coeruleus (LC) consists of a small cluster of noradrenergic neurons in the brainstem. Through widespread axonal connections, the LC is the main source of noradrenaline for most brain structures, including the neocortex and hippocampus (Benarroch, 2018). Release of noradrenaline in the target structures is known to modulate a number of physiological processes including arousal, attention, memory and emotional processing and exerts powerful control over circuit excitability and plasticity (Berridge & Waterhouse, 2003; Harley, 2007; Sara, 2009). In the context of epilepsy, noradrenaline is generally considered an anticonvulsant neuromodulator (Ghasemi & Mehranfard, 2018). Lesions to the LC-noradrenergic system are known to result in severe exacerbation of seizures in rodent epilepsy models (McIntyre et al., 1979; Mishira et al., 1994). Meanwhile, although less consistent, therapeutic interventions elevating noradrenergic transmission are often observed to reduce the severity of seizures (Clinckers et al., 2010; Raedt et al., 2011; Vermoesen et al., 2012). Other studies, however, have claimed proconvulsant effects of noradrenaline under certain conditions, which has been attributed to activation of different subtypes of adrenoceptors of the α- and β-families (Ghasemi & Mehranfard, 2018). The diversity of adrenoceptors throughout various brain areas and in diverse cell populations, including neuronal (Berridge & Waterhouse, 2003) and glial cell populations (Batiuk et al., 2020), may explain the multitude of effects observed. Next to that, the difference in affinity between the different adrenoceptor subtypes, could point towards concentration-dependent effects, depending on a gradual but step-wise recruitment of respectively α_2_-, α_1_- and β-adrenoceptors (Atzori et al., 2016).

Beyond seizure modulating properties of noradrenaline, there is also evidence that the LC-noradrenergic system may play a more direct role in epileptic disorders. There are indications that the transitioning from a healthy to an epileptic brain following a status epilepticus, also known as epileptogenesis, results in changes in expression patterns of adrenoceptors throughout the brain, which may result in a net decrease in the efficiency of noradrenergic transmission (Corcoran & Weiss, 1990a). Evidence also indicates that the LC-noradrenergic system is activated during chemically and electrically evoked seizures in rodents (Jimenez-Rivera & Weiss, 1989; Silveira et al., 2000, 2002; Silveira, Liu, de LaCalle, et al., 1998; Silveira, Liu, Holmes, et al., 1998). Limbic seizures have been associated with increased expression of cFos, a marker for neuronal activation, in LC neurons (Silveira et al., 2000, 2002; Silveira, Liu, de LaCalle, et al., 1998; Silveira, Liu, Holmes, et al., 1998), while a previous report suggests an increase in LC multiunit activity in seizures evoked by electrical stimulation of the amygdala (Jimenez-Rivera & Weiss, 1989). This suggests that understanding how the LC-noradrenergic system is affected by seizures can be key to understanding seizure pathophysiology.

In this study, we investigated changes in LC-unit activity and LC-noradrenergic transmission during acute hippocampal seizures in rodents applying novel neuroscientific research tools such as high-density electrophysiology, optogenetics and noradrenaline biosensing. The identity of LC neurons was additionally confirmed through optogenetic tagging, where LC specific expression of the inhibitory opsin GtACR2 was combined with blue light illumination, allowing LC neuron identification *in vivo* (Lee et al., 2020; Lima et al., 2009). The recently developed GRAB_NE2m_ biosensor (Feng et al., 2019) was used to assess changes in noradrenergic transmission in the hippocampus during seizures with high subsecond temporal precision.

## Results

### Characteristics of evoked hippocampal seizures

Hippocampal seizures were evoked in anesthetized male Sprague Dawley rats with 10s tetanic trains of perforant path stimulation (20 Hz, 0.2 ms bipolar square wave pulses), using stimulation intensities ranging from 300-800 µA. The seizures were initially characterized by repetitive high-amplitude negative spikes in the local field and a general increase in broad band power (**Figure 1** and **supplementary Figure 1**). Seizure duration was very stable within animals when evoking multiple seizures, but was observed to vary between animals. The mean duration of the first seizure evoked in each experimental session across all animals was 13.1±4.8 seconds (mean ± SD; range 5.3-24.0 seconds).

**Figure 1:**
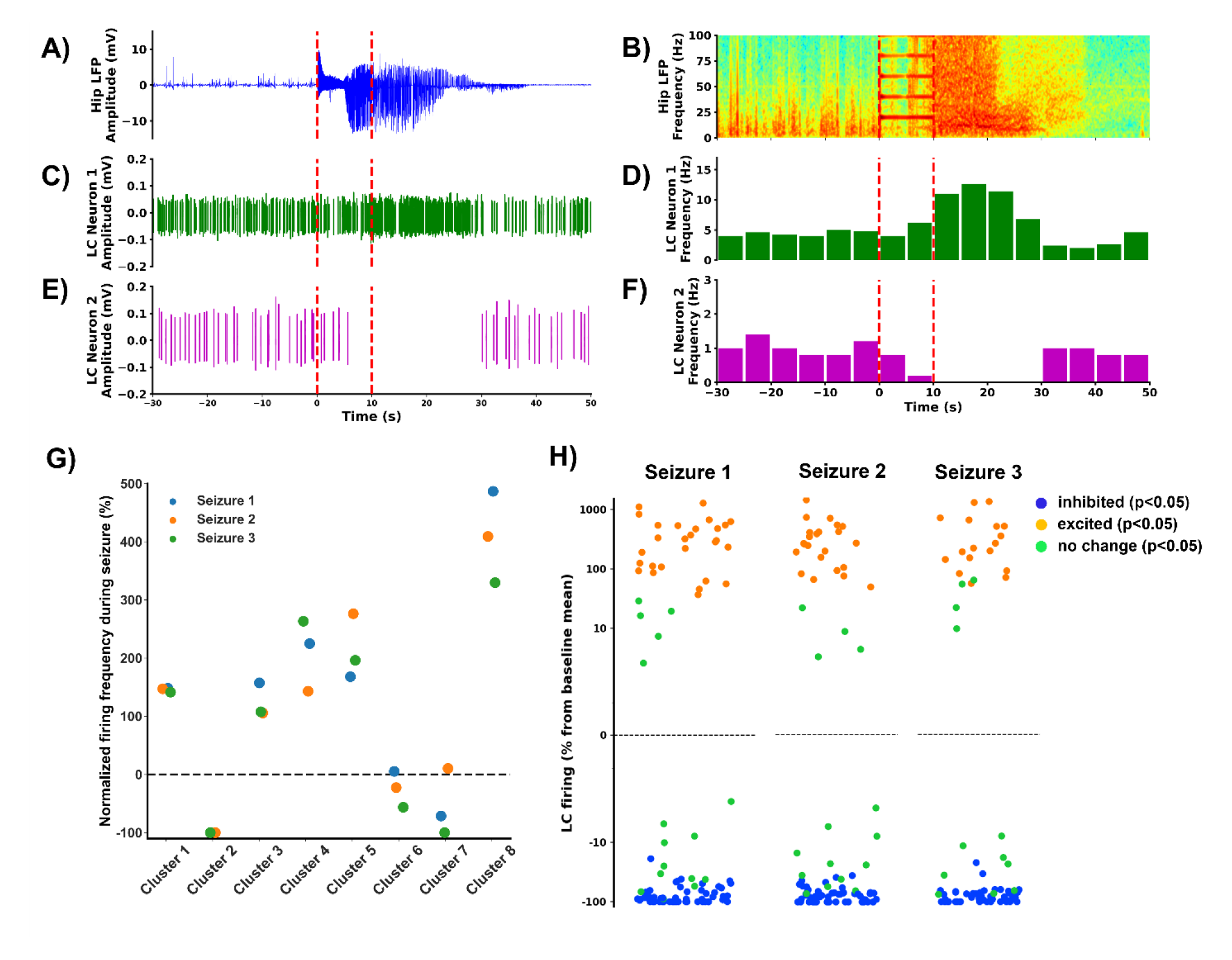
Example of an electrically evoked hippocampal seizure and associated changes in firing rates of locus coeruleus (LC) neurons. Hippocampal seizures were induced with 10s tetanic burst stimuli delivered in the perforant path, denoted with red dashed lines. A) Hippocampal local field potential (LFP) trace, B) a spectrogram of the hippocampal LFP, C) and D) example of LC neurons displaying increased firing during hippocampal seizures, E) and F) example of a LC neurons showing decreased firing during hippocampal seizures, G) normalized changes in firing rates of the LC neurons identified based on single channel tungsten needle recordings and H) normalized changes in firing rates of neurons identified from multi-channel probe recordings. In G) and H), firing rates during hippocampal seizures have been normalized to the mean firing rate the minute before seizure induction.

### Evoked hippocampal seizures modulate the firing of locus coeruleus neurons

LC neurons were targeted using three different electrode set-ups: 1) a single-channel tungsten microelectrode, 2) a 32-channel silicon probe and 3) a 32-channel silicon probe with a Ø100 µm optical fiber attached, terminating at the most proximal electrode contact.

Single channel recordings yielded a total of 8 clusters of neurons recorded in 5 out of 9 animals. Spike sorting based on the single channel recordings did not always allow unambiguous characterization of single-unit activity, for which the putative LC neurons isolated in the single channel LC recordings are referred to as ‘LC neuron clusters’ and may in some cases reflect multiple neurons. All identified neuron clusters responded with a typical burst-inhibition to noxious stimuli and were inhibited following administration of clonidine (0.04 mg/kg, I.P.), an agonist of the α_2_-adrenergic receptor, upon conclusion of the experiments. Evoked hippocampal seizures were associated with profound changes in firing rate of most of the recorded LC neuron clusters, although both increased and decreased firing was observed (**Figure 1**). The first evoked hippocampal seizure resulted in increased firing in five LC neuron clusters, while one LC neuron cluster was significantly inhibited during seizures. Two LC neuron clusters showed no significant changes in firing during seizures. The response of LC neuron clusters was observed to be consistent over consecutively evoked seizures. Additional LC recordings were performed using high-density silicon probes to increase the yield of neurons recorded and the validity of the spike sorting process.

Using a 32-channel high-density electrophysiology probe, LC activity was identified in 9 out of 14 animals. In two animals, recordings were performed both ipsilateral and contralateral to the side of the perforant path stimulation. In total, recordings resulted in the identification of 97 LC neurons that responded to noxious foot pinches or were tagged using optogenetics with LC-specific expression of the inhibitory opsin GtACR2 (**Figure 8**, **supplementary Figure 2**). Identified neurons further responded with pronounced inhibition following administration of clonidine (0.04 mg/kg, I.P.). Before and between seizures, the mean firing rate of recorded neurons was 2.3 Hz ± 1.4 Hz (median 2.1 Hz). In contrast to the single-channel recordings, a majority of neurons recorded, 54 (56%), were significantly inhibited during seizures. Only 27 (28%) LC neurons were excited during seizures, while a minority of 16 (16%) LC neurons showed no significant change in firing rate. Changes in LC firing were typically aligned with the onset of hippocampal seizure activity. The response of individual neurons was observed to be consistent over subsequently evoked seizures, with 59% and 62% of LC neurons being inhibited and 25% and 22% of LC neurons being excited during the second and third seizure, respectively. The responses of individual LC neurons over consecutively evoked seizures were highly correlated (r=0.87, 0.80 and 0.92 for seizure 1 vs. seizure 2, seizure 1 vs. seizure 3 and seizure 2 vs. seizure 3, respectively, **supplementary Figure 3**).

Around 35% of the 97 neurons recorded from the LC, were recorded contralateral to the stimulated perforant path. However, the fraction of neurons responding with excitation vs. inhibition in relation to the evoked seizure for contralateral (44%/52%) vs. ipsilateral (32%/56%) recordings was not found to substantially differ.

### Characteristics of inhibited versus excited locus coeruleus neurons

Next, we aimed to identify potentially separating characteristics of inhibited vs. excited LC neurons. Previous studies have indeed suggested the presence of distinct types of LC neurons that can be distinguished on the basis of spike waveforms and spike rate (Totah et al., 2018a). Our data, however, did not show any tendencies for such clustering (**supplementary Figure 4A-C**). Next we assessed whether spike width, peak asymmetry in spike waveforms, mean spike rate or median interspike interval could predict the response of LC neurons to evoked hippocampal seizures. However, no such relationship was indicated by our analyses (**supplementary Figure 4D-G**).

Additionally, the type of coupling between LC neurons, broad (tens of milliseconds) or sharp (< millisecond) coupling, was assessed to investigate a possible correlation to the excitatory or inhibitory activity during an evoked hippocampal seizure (Totah et al., 2018). Both types of coupling were observed in our data, when looking at the spontaneous spiking patterns of neurons in the absence of seizures, and at rates comparable to those observed in literature (**supplementary Figure 5**). However, the rates of significant cross-correlograms were similar for pairs of inhibited neurons, pairs of excited neurons and pairs of inhibited and excited neurons (**supplementary Figure 5E**). Finally, we evaluated cross-correlation between neurons that were excited during seizures (58 neuron pairs). A limited number of spikes was measured due to a limiting sampling period (50-seconds around the seizures, 2-3 seizures per recording). The fraction of correlated neurons during seizures was similar to the fraction during interictal periods. Although not all neuron pairs with interictal coupling showed coupling during seizures, preictal sharp coupling was nevertheless observed to predict coupling during seizures (p<0.05, Fisher’s exact test).

In a final attempt to understand potential differences between neurons inhibited or excited in relation to seizures, we focused on anatomy. The high-density multi-channel probe enabled us to compute topographies of the changes in multiunit-activity across the probe in relation to seizures or other events. Beyond confirming the presence of both inhibited and excited LC neurons in relation to seizures, such analysis additionally indicated anatomical separation of clusters of neurons responding to seizures with inhibition vs. excitation (**Figure 2**). Although with some degree of variability, such pattern was observed in multiple animals (**supplementary Figure 6**) and was observed to be highly consistent over the course of multiple, consecutively evoked hippocampal seizures (**Figure 2**).

**Figure 2:**
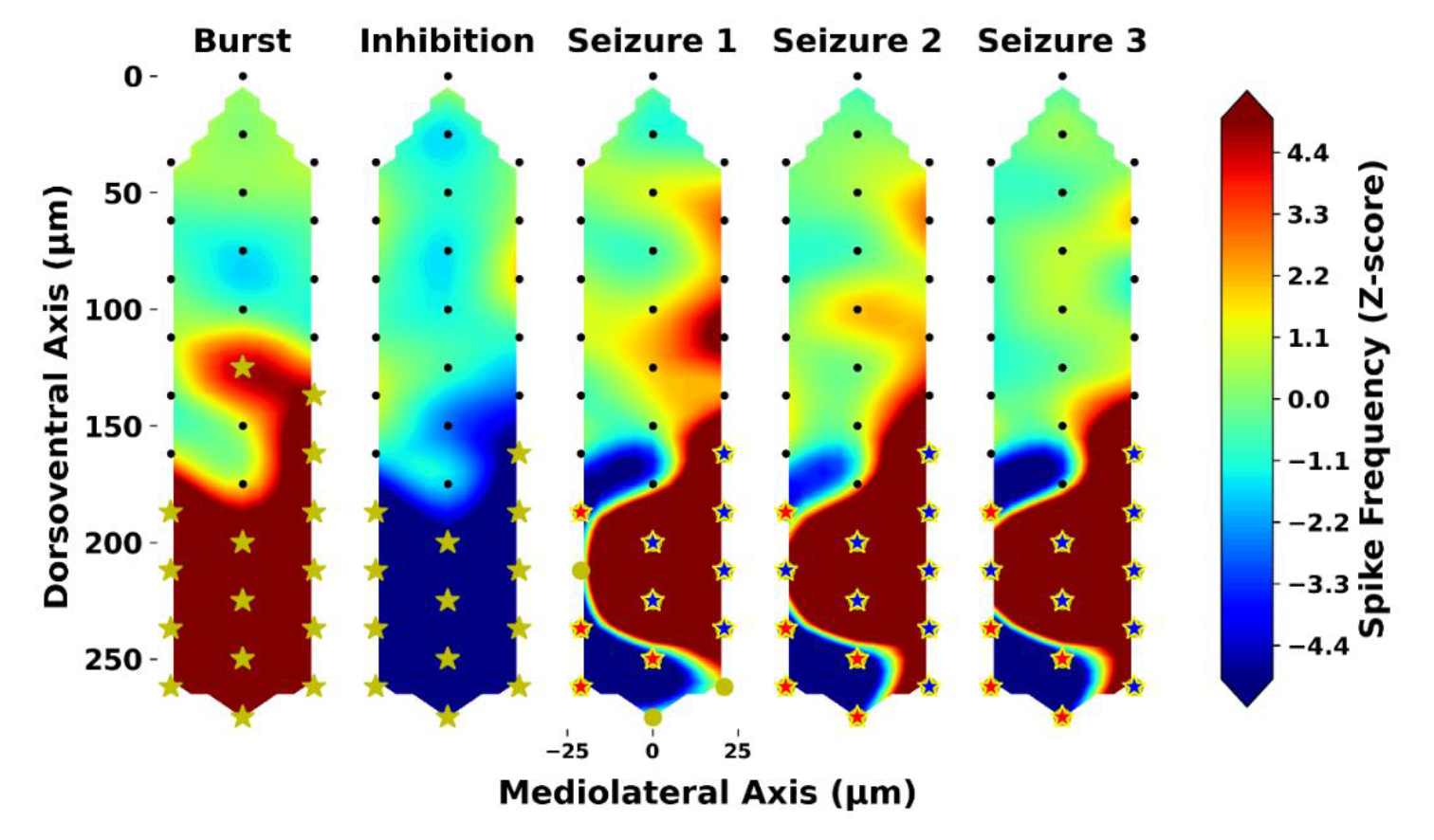
A filled contour plot to illustrate multiunit firing rates across the 32-channel recording probe in relation to specific events. Firing rates have been geometrically interpolated by means of cubic interpolation. Left, in the first two plots, is an illustration of the average noxious foot pinch response characterized by a burst, subsequently followed by inhibition. Significant responses at an individual channel level are indicated with yellow stars. The example provided shows profound and very consistent changes in multiunit activity across the probe in relation to repeated seizure inductions. Significant inhibition is noted with red stars and significant excitation is noted with blue stars. Channels marked as LC channels based on significant burst inhibition in response to a noxious foot pinch are additionally noted with a yellow edge around the stars.

### Coupling of LC neurons to the hippocampal LFP during evoked hippocampal seizures

From initial observations it was clear that changes in LC activity coincided with the occurrence of seizure activity in the hippocampal LFP. In a few cases, intermittent seizure activity in the hippocampal field was even associated with additional bursts of activity in these neurons (**supplementary Figure 9**). A quantitative analysis of this temporal association was performed by means of cross-correlation of 1-second hippocampal LFP amplitude bins and LC neuron spike rate bins (**Figure 3**). Since it was difficult to analyze the hippocampal LFP reliably during seizure induction, due to electrical stimulation artefacts and difficulties dissociating evoked field activity from seizure activity, analysis focused on the 50 seconds immediately following stimulation and thus in particular the timing of seizure termination and normalization of LC firing (**Figure 3A**). Analysis revealed neurons displaying strong anticorrelation and correlation to the amplitude of the hippocampal LFP, reflecting LC neurons showing inhibition and excitation in relation to seizures, respectively.

**Figure 3:**
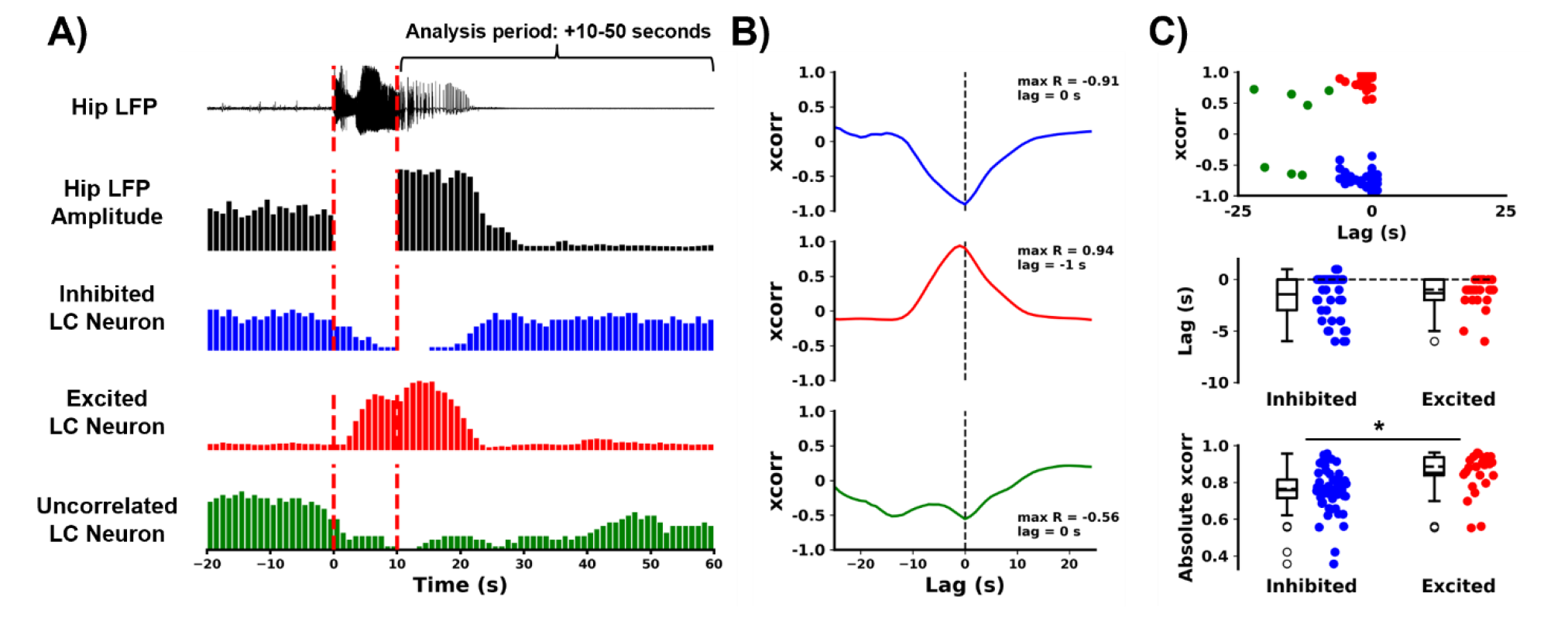
Temporal association between seizure activity in the hippocampal local field potential (LFP) and changes in the firing of locus coeruleus (LC) neurons. A) Cross-correlation analyses were based on 1 second bins of hippocampal LFP amplitude and firing rates of individual LC neurons. B) Cross-correlation (xcorr) analysis revealed the presence of LC neurons showing strong anticorrelation to changes in hippocampal LFP amplitude (inhibited neurons), neurons showing strong direct correlation (excited neurons) and neurons showing weak correlation (uncorrelated neurons) to the 50 seconds immediately following electrical seizure induction. C) Anticorrelated and directly correlated LC neurons could be separated with a density-based clustering analysis which showed similar lag, but a strong cross-correlation of excited vs. inhibited neurons (* denoting statistical significance).

Few neurons showed no clear correlation with the hippocampal LFP (**Figure 3B**). A density-based clustering analysis on the basis of peak cross-correlation and lag indeed allowed the identification of two clear clusters of neurons showing strong anticorrelation (n = 51 neurons) and correlation (n = 25 neurons) to the amplitude of the hippocampal LFP (**Figure 3C**). While there was no detectable difference in the lag of their association, neurons excited by seizures displayed a stronger peak cross-correlation with the amplitude of the hippocampal LFP than inhibited neurons (0.85±0.11 vs. 0.78±0.09, p<0.001). In the vast majority of cases, peak correlation was observed with a negative lag or at 0, indicating that firing of LC neurons normalized before normalization in hippocampal LFP amplitude.

A few excited neurons were additionally observed to display coupling to the phase of the hippocampal LFP (8-16 Hz), which exceeded what could be expected by chance even after Bonferroni correction (**Figure 4**). None of these neurons showed any corresponding coupling to the phase of the hippocampal LFP in the 60 seconds prior to seizure induction (**supplementary Figure 9**). Three of the five neurons displaying significant phase coupling to the hippocampal seizure LFP were identified in the same recording and were observed to generally bind to the throughs of the filtered 8-16 Hz hippocampal LFP signal. During a second seizure, only one of the three neurons retained significant coupling to the hippocampal seizure LFP (p<0.01), while coupling of a second neuron was trending towards significance following Bonferroni correction (p=0.085) (**supplementary Figure 8**).

**Figure 4:**
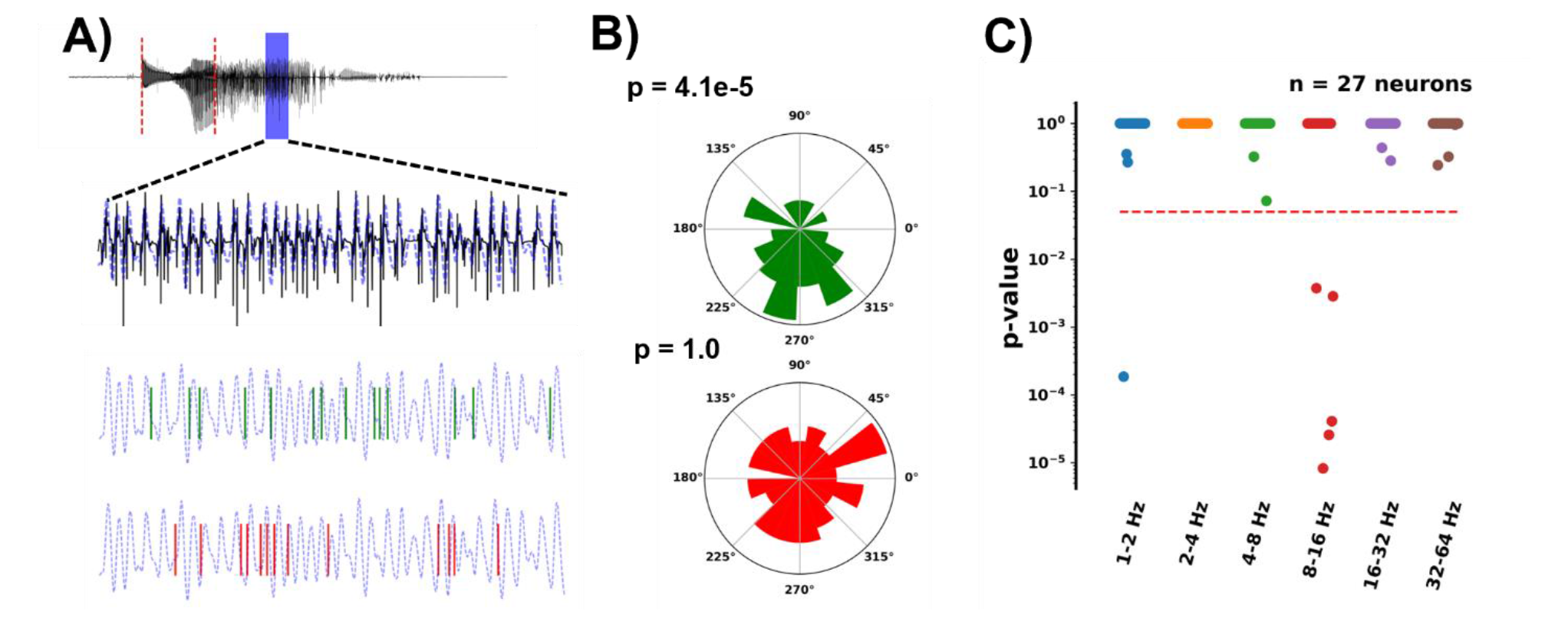
Phase preference of locus coeruleus (LC) neurons to the hippocampal local field potential (LFP) during evoked seizures. A) Example of hippocampal LFP, which was filtered in varying frequency bands (blue dashed lines) and 1. B) example of an LC neuron displaying significant phase preference to the 8-16 Hz bandpass filtered hippocampal LFP (green) and an uncoupled neuron (red). C) An overview of all neurons following significance correction, showing that 5 LC neurons displayed significant coupling to the hippocampal 8-16 Hz LFP.

An additional analysis assessed coupling between spiking of LC neurons and the occurrence of spikes in the hippocampal LFP in relation to the seizures (**supplementary Figure 9**). While two neurons showed statistically significant increased likelihood of spiking around the time of hippocampal LFP spikes, the relationship did not remain following Bonferroni correction for multiple testing.

### Photometric quantification of GRAB_NE2m_ in relation to hippocampal seizures

Beyond assessing changes in LC unit activity, a fluorescent sensor for noradrenaline, GRAB_NE2m_, was used to quantify changes in noradrenaline release in the hippocampus in relation to evoked hippocampal seizures (**Figure 5**). Induction of hippocampal seizures was observed to consistently increase GRAB_NE2m_ fluorescence both in anesthetized (31.5 ± 33.8%) as well as awake (23.9 ± 16.2%) conditions, indicating an increase in extracellular noradrenaline concentration. While the extent of the change in GRAB_NE2m_ fluorescence was highly variable at a group level (range anaesthesia: 5.6 - 107.1%, range awake: 6.9 – 60.1%, **Figure 5D** **and** **H**), the peak changes observed were nevertheless consistently far beyond what could be expected by chance for individual animals (z-score range anaesthesia: 6.8 – 502.2, z-score range awake: 16.0 - 380.6, **Figure 5E** **and** **I**). Similar to previous observations of temporal correlations between changes in hippocampal LFP amplitude and LC unit activity, a strong correlation was observed between changes in GRAB_NE2m_ fluorescence and hippocampal LFP amplitude in relation to seizures. Negative lags indicate a tendency for the GRAB-signal to peak prior to normalization of the hippocampal LFP.

**Figure 5:**
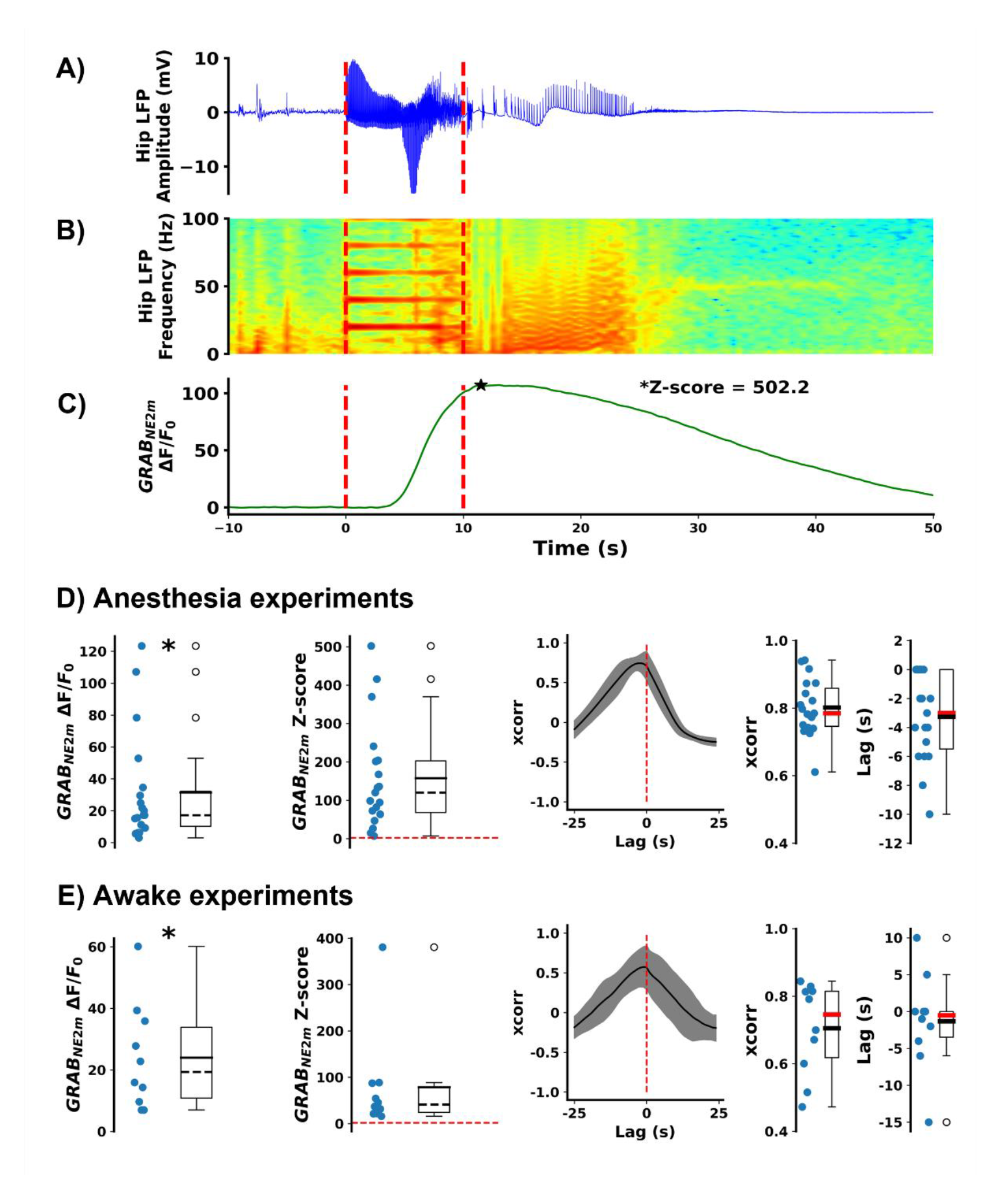
Changes in noradrenergic transmission in the hippocampus were assessed using the fluorescent sensor for noradrenaline GRAB_NE2m_ in combination with fiber photometry. A-C) Perforant path stimulation induced seizures (red dashed lines) were associated with an increase in GRAB_NE2m_ fluorescence. Similar outcomes were observed in anesthetized (D) and awake rats (E), * indicates the peak response in fluorescence for one animal after a seizure. Changes were observed to be highly significant both at a group level and individual level. Changes in GRAB_NE2m_ fluorescence were further found to be correlated to changes in hippocampal LFP amplitude when cross correlating 1-second amplitude bins over the 50-second period following electrical stimulation.

## Discussion

Previous studies have indicated that various types of seizures in rodent seizure models modulate the LC-noradrenergic system (Jimenez-Rivera & Weiss, 1989; Silveira et al., 2000, 2002; Silveira, Liu, de LaCalle, et al., 1998; Silveira, Liu, Holmes, et al., 1998). In this study, we used a 32-channel high-density probe for LC unit recording and the recently developed GRAB_NE2m_ biosensor for noradrenaline detection, to perform a detailed assessment of the effect of evoked hippocampal seizures on LC-noradrenergic transmission. We found that hippocampal seizures consistently modulated LC activity and led to a strong release of noradrenaline in the hippocampus. Understanding the physiological or pathophysiological implications of this release is likely to be important to understanding the pathophysiology of seizures.

In a series of initial single channel recordings of LC unit activity with tungsten electrodes, we recorded few neuron clusters of which a majority showed increased firing during seizures (5/8), which thus aligned with our initial expectations (Jimenez-Rivera & Weiss, 1989; Silveira, Liu, de LaCalle, et al., 1998; Silveira, Liu, Holmes, et al., 1998). Subsequent high-density recordings using 32-channel silicon probes, yielded a higher number of LC neurons (97). However, here, most neurons (∼55%) were inhibited in relation to seizures, while only a small subset of neurons (∼25%) increased their firing. The responses of neurons were consistent over consecutively evoked seizures. Interestingly, literature does indeed indicate distinct classes of LC neurons (Totah et al., 2018). In particular, evidence suggests the presence of slow spiking LC neurons with broad waveforms located predominantly in the dorsal LC, while the ventral LC contains neurons characterized by faster spike rates and narrow waveforms (Totah et al., 2018). Our data, however, did not show any clear clusters of neurons based on spike wave forms and firing rate, but showed the presence of one rather uniform cluster with a mean firing rate of around 2 Hz, which is on the high side relative to typical reports in literature. One explanation, although we have no conclusive evidence hereof, is that we likely recorded mainly from the dorsal LC, as recordings were usually initiated upon the first encounter of LC activity, without lowering the electrode or probe further with the risk of losing LC activity. Here, it is also important to note that LC recordings were conducted using state-of-the-art methodology, including optogenetic tagging of LC activity, resulting in a high level of confidence that LC neuronal activity was being recorded.

The lack of distinct classes of LC neurons in our data further meant that any attempts to separate or characterize inhibited and excited LC neurons on the basis of the baseline firing frequency or waveforms, did not show any meaningful correlations. Instead, we calculated cross-correlation of neurons, which has been described to a limited extent in literature and which is thought to reflect either common synaptic inputs (coupling over tens or hundreds of milliseconds) or electrotonic coupling via gap junctions (coupling within a millisecond) (Totah et al., 2018). Indeed, we observed both types of coupling in our data, and at rates comparable to previous reports (Breton-Provencher et al., 2021). However, although we initially hypothesized a higher rate of coupling among excited neuron pairs as a potential explanation for common excitatory drive, this did not appear to be the case in our data.

Thus, in a final attempt to comprehend possible differences between neurons that respond differently to seizures, we computed topographies of multi-unit activity across the 32-channel probes used for recordings. In this case in at least half of the recordings (5/9), anatomical separation of inhibited vs. excited activity was observed. Interestingly, and contrary to early evidence (Berridge & Waterhouse, 2003), more recent studies suggest that the LC is not a single uniform unit, but that LC neurons are arranged in anatomically distinct modules in accordance with their efferent projection patterns (Chandler et al., 2019; Uematsu et al., 2017). Considering that we consistently observed an increase in noradrenergic transmission in the hippocampus, as shown in the GRAB_NE2m_ experiments, at least part of the neurons excited in relation to hippocampal seizures must be neurons projecting to the hippocampal formation. A definite proof hereof, however, would require either antidromic tagging or the use of e.g. retrogradely transduced opsins (Li et al., 2016). Secondly, it must be considered that a majority of LC neurons (56%) recorded were still inhibited during seizures. With the proposed modular arrangement of the LC in mind (Chandler et al., 2019; Uematsu et al., 2017), it is therefore likely that inhibited neurons innervate other targets than the hippocampal formation, which therefore may result in a drop in noradrenaline concentrations in other parts of the central nervous system in relation to seizures.

Exactly how evoked hippocampal seizures result in modulation of LC neurons, and strong release of hippocampal noradrenaline, remains unclear. Considering that seizures were evoked by electrical stimulation of the perforant path, which is located externally to the hippocampus, it is unlikely that LC projections were stimulated directly and activated antidromically. Changes in LC firing were further aligned with the onset of seizure activity in the hippocampus, which typically had a 5-7 second latency from the onset of the electrical stimulation, further countering this argument. Although the LC provides noradrenergic innervation to the hippocampus (Mason & Fibiger, 1979), there are no known direct projections from the hippocampus to the LC (Sara & Bouret, 2012). Rather, pathways to the LC from the hippocampal formation are polysynaptic connections, such as those via the amygdala or the prefrontal cortex (Breton-Provencher et al., 2021). Afferent projections from the prefrontal cortex to the LC are ionotropic glutamatergic connections, while projections from the amygdala release corticotropin releasing hormone (CRH), which induces excitation through activation of G-protein coupled receptors (Breton-Provencher et al., 2021). This motivated a closer look at potential correlations between the hippocampal local field potential and LC unit activity in relation to seizures. Here, it was clear that changes in LC activity in most cases were confined specifically to the period of the evoked hippocampal seizure. The fast onset and normalization of LC activity following termination of seizure activity would not suggest the involvement of metabotropic receptors which usually would have slow but long-lasting effects. Rather, the data suggests a strong excitatory input to LC neurons excited by hippocampal seizures via a yet undefined ionotropic glutamatergic polysynaptic pathway. What further strengthens this hypothesis is the observation that some of the neurons, observed to be excited in relation to seizures, showed significant phase coupling to the hippocampal LFP during the seizure.

The mechanism via which a majority of neurons are inhibited, on the other hand, may be a result of auto- or hetero-inhibition. LC neurons are known to presynaptically express inhibitory α_2_-adrenoceptors as autoreceptors, both presynaptically at the axon terminals, but also on the cell bodies and dendrites (Berridge & Waterhouse, 2003). LC neurons influence the activity of surrounding LC neurons through somatic and dendritic release of noradrenaline (Andrade & Aghajanian, 1984; Huang et al., 2012). Strong excitation of a subset of neurons, as observed in the present study, may thus result in inhibition of surrounding LC neurons.

The key takeaway from this study, with potentially strong clinical implications, is that evoked hippocampal seizures in rodents consistently result in profound release of noradrenaline during seizures. Importantly, we provide evidence of this not only in anesthetized rats, but also in freely moving awake rats. Although, at present the extent to which the observed phenomenon is generalizable to other types of seizures is unknown, there are a number of observations that could indicate that a profound release of noradrenaline is at least a generalizable mechanism of limbic seizures. For example, we previously observed that evoking a status epilepticus by infusing pilocarpine into the hippocampus in awake rats led to a strong increase in concentrations of noradrenaline in the hippocampal extracellular space, measured with microdialysis (Raedt et al., 2011). Another study showed that limbic seizures, evoked by electrical stimulation of the amygdala, resulted in an increase in LC multiunit activity in awake rats and increased levels of noradrenaline in the LC and amygdala. Silveira *et al.* further published a series of studies showing increased cFos expression in LC neurons following acute limbic seizures in rats evoked with amygdala kindling, kainic acid or flurothyl (Silveira et al., 2000, 2002; Silveira, Liu, de LaCalle, et al., 1998; Silveira, Liu, Holmes, et al., 1998). Further studies should address the extent to which e.g. neocortical seizures and partial vs. generalized seizures similarly modulate LC-noradrenergic transmission.

Noradrenaline is a powerful neuromodulator with profound effects on a number of physiological (and pathophysiological) processes. Considering that the study at hand shows a consistent and profound release of noradrenaline during hippocampal evoked seizures, addressing the physiological and/or pathophysiological implications of this response may be necessary to understand the pathophysiology of seizures. One of the most straight forward deductions is the role of noradrenaline in modulating arousal and behavioral states (Berridge & Waterhouse, 2003). In general, levels of noradrenaline are thought to correlate to levels of alertness. It is therefore puzzling that limbic seizures typically are associated with impairment or even loss of consciousness (Englot et al., 2010). While we indeed consistently saw an increased release of noradrenaline in the hippocampus, it is very possible that noradrenergic transmission decreases in other areas relevant to consciousness (Gummadavelli et al., 2015) or that other mechanisms than release of noradrenaline in relation to seizure mediate impaired consciousness associated with limbic seizures.

Noradrenaline is further known to affect physiology relevant to seizures, such as excitability and plasticity. Although noradrenaline is predominantly seen as an anticonvulsant neuromodulator (Ghasemi & Mehranfard, 2018), this is typically observed under conditions that involve a low-moderate facilitation of LC activity and is predominantly (Corcoran & Weiss, 1990a) associated with activation of α-adrenoceptors, which have relatively higher affinity of noradrenaline than β-adrenoceptors (Atzori et al., 2016). High levels of noradrenaline, which may to a larger extent activate β-adrenoceptors has been associated with enhanced excitability in hippocampal circuits (Harley & Milway, 1986; Heginbotham & Dunwiddie, 1991) and a mixture of both anti- and pro-convulsant effects in rodent seizure models (Ghasemi & Mehranfard, 2018). The question thus remains whether release of high levels of noradrenaline in relation to seizures protect against or facilitate seizure activity.

Beyond immediate effects of noradrenaline on excitability, release of noradrenaline is known to enhance plasticity, similarly through activation of β-adrenoceptors. For example, when high frequency stimulation is paired with noradrenaline, induction of long-term potentiation (LTP) is enhanced (Hansen & Manahan-Vaughan, 2015). Similarly, the consolidation of fear conditioning (Seo et al., 2021) or spatial memory (Hansen & Manahan-Vaughan, 2015) is enhanced when paired with noradrenaline release. A consequence of noradrenaline release in relation of seizure activity may be enhanced consolidation of pathological seizure activity and facilitated formation of seizure networks, which could play a role in epileptogenesis. This hypothesis has partially been studied in previous kindling experiments, by applying lesions to the LC-noradrenergic system (Corcoran & Weiss, 1990). Such lesions, however, are associated with near complete and chronic depletions of noradrenaline in the brain, which strongly exacerbates baseline seizure severity (Giorgi et al., 2003), which severely complicates the interpretation of kindling experiments. Addressing the role of noradrenaline released in relation to seizures will instead require tools that allow LC specific modulation with high temporal precision and with sufficient potency to counter the strong excitatory input to the LC during seizures.

In this study, we demonstrate that acute electrically evoked hippocampal seizures are associated with strong changes in LC unit activity and strong and consistent time-locked release of noradrenaline. Understanding the role of mass release of noradrenaline during hippocampal seizures is likely to be important to understand seizure pathophysiology.

## Materials and methods

All experiments were conducted with male Sprague Dawley rats (Envigo, The Netherlands, 300-400g at the time of the experiment). Animals were treated according to European guidelines (directive 2010/63/EU). All experiments described in the following were approved by the Animal Experimental Ethical Committee of Ghent University (ECD 19/42 and 22/54). Rats were housed under environmentally controlled conditions (humidity: 40-60%, temperature: 20-23°C), a controlled 12/12h light/dark cycle and food (Rats and Mice Maintenance, Carfil, Belgium) and water *ad libitum*.

### Injection of viral vector in LC

Prior to subsequent experimentation, 6 out of 14 rats used for LC unit recordings were injected in the LC with a viral vector for LC specific expression of the blue-light gated chloride channel GtACR2 (**supplementary Figure 2**). Injection of a CAV2 viral vector preceded acute LC unit recordings by at least 3 weeks. Rats were anaesthetized using a mixture of isoflurane (5% for induction/2% for maintenance, Isoflo, Zoetis, UK) and medical oxygen and fixed in a stereotactic frame. Bregma was lowered 2 mm relative to lambda to inject at an angle of approximately 15 degrees, to avoid the transverse sinus. The LC was subsequently targeted bilaterally with stereotactic coordinates 3.9 mm posterior and 1.2 mm lateral to lambda. At 3 depths, -5.5, -5.8 and -6.1 mm relative to the brain surface, 0.6 µL of the viral vector construct CAV2-PRS-GtACR2-fRED (titer 0.56x10^12^ pp/mL, IGMM, France) was injected at a rate of 100 nl/min, using a 5 µL Hamilton Neuros Syringe (33 gauge, point style 3, Hamilton company, USA) and a Quintessential Stereotaxic Injection System (Stoelting, USA). Following each injection, the needle was left in place for 5 minutes to prevent backflow of the vector. At the end of the procedure, animals were injected with Meloxicam (1 mg/kg, Boehringer Ingelheim, Germany) and left to recover for at least 3 weeks to allow expression of GtACR2.

### LC unit recordings and evoking of hippocampal seizures

Acute LC unit recordings were performed in rats under isoflurane anesthesia (5% for induction/2% for maintenance). Before targeting the LC, a polyimide-coated stainless steel wire electrode (ø70 µm, California Fine Wire, USA) was implanted in the hilus of the dentate gyrus in the left dorsal hippocampus (3.8 mm posterior, 1.9 mm lateral relative to bregma, ∼-3 mm ventral to brain surface) for registration of local field potentials (LFPs). A custom-made bipolar electrode for stimulation, made from two twisted polyimide coated stainless steel wires (ø110 µm, Science products, Germany), was implanted in the left perforant path (3.9 mm lateral to lambda, ∼-2.5 mm ventral to brain surface). Electrodes were adjusted to obtain maximal amplitude of evoked potentials in response to 200-400 mA bipolar square wave pules (0.2 ms pulse width) delivered to the perforant path. Once in place, these electrodes were sealed in place with dental cement and bound to surrounding bone screws (ø 0.8 mm, Plastics One). An additional wired bone screw placed in the right frontal bone was used for grounding and as a reference for all electrophysiological recordings in these experiments.

The LC was subsequently targeted using three different electrode set-ups: 1) a single-channel tungsten microelectrode (0.00800/200 µm shank diameter, recording impedance ≥1.5 MΩ, FHC, USA), (n=9 rats), 2) a 32-channel silicon probe, model A1x32-Poly3-10mm-50-177, recording impedance ≥1.0 MΩ, Neuronexus, USA), (n=8 rats), 3) a 32-channel silicon probe (model A1x32-Poly3-10mm-25-177, recording impedance ≥1.0 MΩ, Neuronexus, USA) with a Ø100 µm optic fiber (0.39 NA, FT100EMT, Thorlabs, Germany) attached terminating at the most proximal electrode (n=6 rats). LC recordings with the high-density silicon probes were performed to increase the yield of neurons recorded and the validity of the spike sorting process, which is easier when action potentials of single neurons can be recorded over multiple electrode contacts in close proximity. The latter probe was used for LC recordings in rats injected with viral vector for expression of GtACR2, to additionally confirm the identity of LC neurons.

The initial LC coordinates used were 3.9 mm posterior and 1.15 mm lateral to lambda at an angle of 15 degrees towards the front. LC activity was usually found 5.5-6.2 mm ventral to brain surface. To confirm the identity of LC activity, different parameters were used: 1) slow, regular firing patterns (1-6 Hz), with broad spike waveforms (∼1 ms), 2) a burst response (10-15 Hz) followed by inhibition (300-700 ms) upon application of a noxious foot pinch (Vazey et al., 2018) and 3) optogenetic identification of LC activity. For subsequent offline identification of LC activity, multiple trials of noxious foot pinch stimuli were performed (typically around 10) separated by at least 30 seconds. For animals expressing GtACR2 in the LC, a number of light trials (typically around 10) were performed for additional optogenetic confirmation of LC identity of the recorded neurons. These trials, separated by at least 60 seconds, consisted of 3-7 second light pulses delivered at 4 mW through the Ø100 µm optic fiber probe attached. Upon registration and stabilization of LC activity (typically waited ∼15 minutes), the experimental protocol was initiated.

Subsequently, a number of hippocampal seizures were evoked with 10 second tetanic trains of electrical stimulation (20 Hz, 0.2 ms bipolar square wave pulses, DS4 Bi-Phasic Stimulator, Digitimer, USA). The intensity was gradually increased until a reliable >10 s seizure was evoked in the hippocampal field (typically at an intensity of 300-600 µA). Seizures were separated by at least 10 minutes. The number of seizures evoked varied among animals (minimum was two, maximum was 10) depending on the stability of the LC unit recordings. Recordings, except those including optogenetics, were concluded with an intraperitoneal injection of the α_2_-adrenoceptor agonist clonidine (0.04 mg/kg), to identify noradrenergic neurons that respond with pronounced inhibition or silencing to the injection(Svensson et al., 1975).

### GRAB_NE2m_ photometry experiments

In a sequential experiment, changes in hippocampal noradrenergic transmission, during acute hippocampal seizures, were assessed in 19 Sprague-Dawley rats using the GRAB_NE2m_ fluorescent biosensor for noradrenaline (Feng et al., 2019b). The GRAB_NE2m_ sensor is a modified α_2_-adrenoceptor with green fluorescence protein (GFP) domain which increases its brightness upon binding of NA. Expression of the GRAB_NE2m_ sensor was induced in hippocampal neurons through injection of 600 nl of the viral vector construct AAV9-hSyn-NE2m-mRuby3 (titer 10^13^ pp/mL, WZ Biosciences, USA) at 3.8 mm posterior and 1.9 mm lateral to bregma and at a depth of -3.0 mm relative to brain surface. After at least two weeks, allowing the GRAB_NE2m_ sensor to express, acute experiments were performed under isoflurane anesthesia (5% induction, 2% maintenance). A bipolar stainless steel wire electrode was implanted at the lambda anterior-posterior coordinate and 3.9 mm lateral to lambda at a depth of ∼-2.5-3.0 mm relative to brain surface for perforant path stimulation. A bipolar stainless steel wire electrode was implanted at 5.3 mm posterior and 3.2 mm lateral to bregma and ∼3 mm ventral to brain surface for acquisition of hippocampal LFPs. GRAB_NE2m_ fluorescence was assessed by means of a Ø200 µm optic fiber (0.39 NA, FT200EMT, Thorlabs, Germany) and photometry using the PyPhotometry acquisition board with the time division setting at a 130 Hz sample rate (Akam & Walton, 2019). This system was paired with a 470 nm LED (Doric, Canada), a filter cube (Minicube, Doric, Canada) allowing separation of light used for excitation (470 nm) and emitted fluorescence (∼550 nm), and a Femtowatt Silicon Photoreceiver (Model 2151, Doric, Canada). As in previous experiments, seizures were evoked by 10 second tetanic trains of electrical perforant path stimulation (20 Hz, 0.2 ms bipolar square wave pulses) at an intensity of 300-600 µA, while continuously monitoring GRAB_NE2m_ fluorescence. Seizures were separated by at least 10 minutes. Ten out of 19 animals were implanted for chronic experimentation and allowed to recover for awake experimentation. In these cases, at least a week after the acute experiment, rats were attached to a custom-made electrophysiology-photometry setup allowing free movement via optical (Pigtailed fiber optic rotary joint, Doric, Canada) and electrical commutators. As under anaesthesia, seizures were evoked with 10 second tetanic trains of electrical perforant path stimulation (20 Hz, 0.2 ms bipolar square wave pulses) while recording hippocampal LFPs and GRAB_NE2m_ fluorescence.

### Electrophysiology hardware settings

Hippocampal LFPs were filtered and amplified (1-1000 Hz, 100x or 200x amplification) and digitized at 2 kHz with a 16 bit resolution over a ±10V input range using a National Instruments data acquisition device (National Instruments, Austin Texas, USA). LC unit activity acquired with a single channel tungsten needle electrode was filtered and amplified (300-3000 Hz, 10000x amplification) and digitized at 30 kHz with a 16 bit resolution over a ±10V input range using acquisition hardware (Micro1401-4) from Cambridge Electronic Design (Cambridge, UK). LC unit activity acquired using a 32-channel silicon probe from Neuronexus was analog filtered (1-7000 Hz) and digitized at 30 kHz with a 16 bit resolution over an effective ±6.4 mV input range using Intan amplifier boards (Intan Technologies, Los Angeles, California, USA). Data acquisition was done using an Open Ephys acquisition board (www.open-ephys.org) and stored for offline analysis.

### Data analysis and statistical analysis

All digital signal processing, analysis and visualization was performed offline using custom scripts or software made in Python. Spike-sorting of single-channel LC spike-data was done using Spike2 (Cambridge Electronic Design, Cambridge, UK). Spike sorting on the basis of multi-channel recordings was done using SpyKING-Circus (version 1.06) (Yger et al., 2018). Manual curation and merging of isolated spike clusters was performed using the Phy GUI, from which spike trains were extracted for the respective putative single neurons.

#### Identification of LC activity

LC activity was identified in 9 out of 14 animals, on the basis of a well characterized burst-inhibition response to noxious stimuli (foot pinches), with optogenetic tagging, where possible, or as a response to the α_2_- adrenoceptor agonist clonidine (0.04 mg/kg, I.P.). During post hoc analysis foot pinch trials were evaluated by binning spike rates in 1 second bins followed by averaging all trials. Z-scored bins were then computed based on mean and standard deviation of the baseline, defined as 20 seconds prior to stimulus application. A significant burst-inhibition response to a noxious foot pinch was defined as an initial burst bin exceeding a Z-score of 3 (p<0.01), followed by an inhibition bin of less than -3 (p<0.01; for example, see **Figure 7A** **and** **B**). Light trials were binned using a bin size equal to the duration of the light stimulus (3-7 seconds), followed by averaging of trials. Z-scoring was done based on the mean and standard deviation of all bins around the light stimulus (for example, see **Figure 7C** **and** **D**).

**Figure 7:**
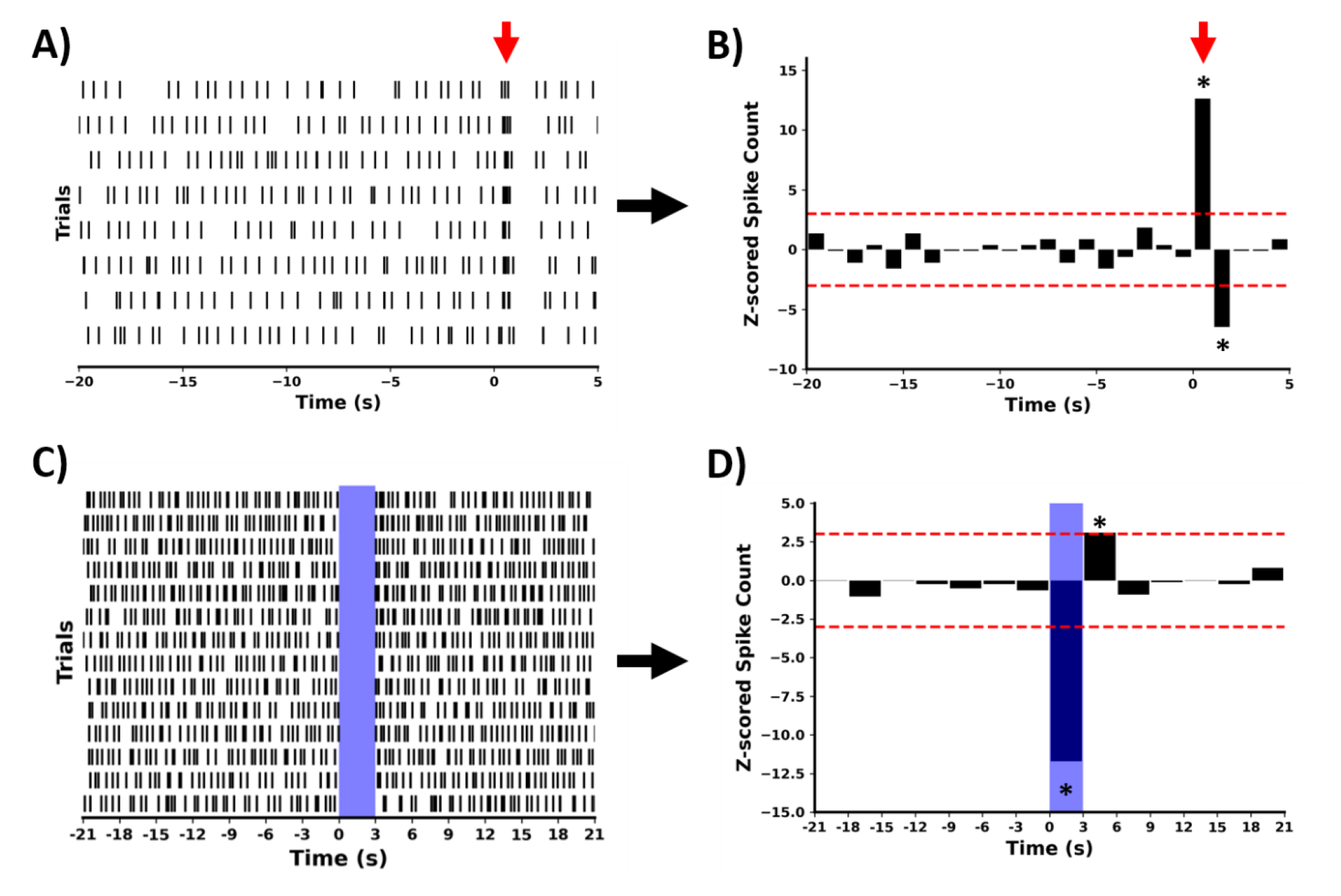
Identification of LC neurons on the basis of a response to a noxious mechanical foot pinch stimulus (A and B), or optogenetic tagging with GtACR2 (C and D). Trials were binned and averaged and a Z-score was calculated on the basis of the activity recorded before and after the application of the stimulus. A significant response to a noxious foot pinch or light (indicated with an asterisk here) was defined as a Z-score of ± 3 standard deviations (threshold indicated with horizontal dashed red lines). Application of the noxious mechanical foot pinch is indicated with a red arrow in A) and B), while blue light application in C) and D) is indicated with blue shading.

#### Evaluation of LC neuronal activity during hippocampal seizures

The activity of identified LC neurons during evoked hippocampal seizures was evaluated by initially computing the spike rate in 5 second bins followed by Z-scoring derived from the mean and standard deviation of the 300 seconds preceding the evoked seizure. In two cases, both within the same recording, the evoked seizures did not last for 10 seconds and these seizures were thus excluded from further analysis. A significant response to seizures was defined as a deviation of more than 1.96 standard deviations (p<0.05). The consistency of responses of LC neurons over consecutive seizures was evaluated using Pearson correlation analysis.

#### Characteristics of LC neurons

Characteristics of LC neurons were evaluated by three metrics: average spike rate, median interspike interval and waveform asymmetry. The average spike rate and median spike interval were computed using data from the entire recording, excluding a 60 second period around evoked seizures and excluding any period post clonidine injection. Over the same period, the spike waveforms of the neurons were extracted from the electrode contact where the neuron displayed the highest amplitude. Waveform asymmetry was computed as an index of the amplitude of the two positive peaks of the spike waveforms as suggested by Totah *et al.* (Totah et al., 2018a).

To look at cross-correlation of neurons in the period between seizures (inter-ictal activity), spikes from the same time-frame between seizures were selected. Two types of cross-correlation common to LC neurons were assessed as described by Totah *et al.* (Totah et al., 2018a). A ‘sharp’ type of coupling, hypothesized to reflect electrotonic coupling of neurons, was calculated over a window of ±3 ms with a bin size of 0.5 ms. A ‘broad’ type of coupling, hypothesized to reflect putative common synaptic inputs, was calculated over a window of ±200 ms with a bin size of 10 ms. For statistics, 200 jittered surrogate cross-correlograms were computed per spike pair by jittering the spikes with ±1 ms (sharp coupling) or ±200 ms (broad coupling). Two significant thresholds were computed: 1) a local ‘per-bin’ threshold computed from the mean ±3 standard deviation of the jittered cross-correlograms, and 2) a global ‘per cross-correlogram’ threshold calculated as the 99% minimum and maximum at any points of the computed jittered cross-correlograms. Cross-correlograms with bins exceeding both of these thresholds were considered ‘significant’. Rates of significant cross-correlograms between conditions were compared in Fisher’s exact tests.

#### Topographies of multiunit activity

The topography of multiunit activity, was computed by extracting threshold crossings per electrode contact from the 32-channel silicon probe recordings. Threshold crossings were extracted using SpyKINGCircus (v. 1.06) using a threshold of 7 median absolute deviations from the median of the signals. Per contact, spike rates were binned (1 second bins) and Z-scored based on the mean and standard deviation of firing. Significance was inferred for Z-scores higher or lower than 3 (p<0.01). Topography maps were smoothed using cubic interpolation for graphical representation.

#### Analysis of coupling of LC neuron activity to the hippocampal local field potential in relation to evoked seizures

Temporal association between changes in firing of LC neurons and changes in the hippocampal LFP amplitude were assessed using 50 seconds of data immediately following perforant path stimulation. Hippocampal LFP ‘total’ amplitude was calculated using the Fast Fourier Transform and binning per second. The spike rate of LC neurons was similarly binned per second and cross-correlated with the hippocampal LFP amplitude to assess peak cross-correlation indices. The peak cross-correlation and lag at peak cross-correlation was used for a density-based cluster analysis to separate classes of neurons which subsequently were compared based on the absolute peak cross-correlation and lag at peak cross-correlation using t-tests.

Coupling of LC neurons to the phase of frequency components of the hippocampal LFP, was computed using exponentially scaling zero-phase second order bandpass filters (bands: 1-2 Hz, 4-8 Hz, 8-16 Hz, 16-32 Hz, 32-64 Hz). The instant phase angles of the filtered hippocampal LFP were computed using the Hilbert Transform. The phase angle probabilities of individual LC neurons were statistically assessed using circular statistics by comparing the phase angle distribution to a uniform distribution, using a Rayleight test. Computed p-values were adjusted for multiple comparison using Bonferroni correction.

Similarly, putative coupling of LC spikes to spikes in the hippocampal LFP was assessed. Negative spikes were detected in the hippocampal LFP using a zero-phase highpass filter with a cutoff at 200 Hz followed by the find_peaks function in the Python scipy.signal module. The LC spike count distribution around population spikes was then calculated for a 1-second window around the population spikes and compared to a uniform distribution using a Kolmogorov-Smirnov Test. Computed p-values were adjusted for multiple comparison using Bonferroni correction.

#### Analysis of GRAB_NE2m_ fluorescence data

Photometric data of GRAB_NE2m_ fluorescence acquired in relation to evoked hippocampal seizures was subjected to zero-phase low-pass filtering (cut-off at 1 Hz). The data was subsequently z-scored to the 300 seconds period before seizure induction. Significant changes in GRAB_NE2m_ fluorescence were defined as changes of at least 1.96 standard deviations (p<0.05) from pre-seizure levels. We further assessed changes in fluorescence using the ΔF/F_0_ metric, which were then used to assess changes in fluorescence at a group level using a one-sample t-test. Changes in GRAB_NE2m_ fluorescence were additionally correlated to changes in hippocampal LFP amplitude using one-second bins and cross-correlation.

### Histology

Immediately following all acute recordings, animals were euthanized with an overdose of sodium pentobarbital (Dolethal, 200 mg/kg, intraperitoneal, Vetoquinol, UK), followed by transcardial perfusion with phosphate buffered saline (PBS) and paraformaldehyde (4%, pH 7.4). The brains were isolated from the skull, post-fixed in 4% paraformaldehyde for 24h and cryoprotected for 3-4 days in a sucrose solution of 30% at 4°C. Subsequently the brains were snap-frozen in isopentane cooled in liquid nitrogen. Coronal sections of 40 μm were made using a cryostat (Leica, Germany). For immunohistochemistry, in order to visualize GtACR2 or GRAB_NE2m_ expression, free floating slices were initially rinsed in distilled water, then incubated in 0.5% H_2_O_2_ for 30 min and in 1% H_2_O_2_ for 60 min, to block endogenous peroxidase activity. Subsequently the sections were washed in PBS (2x 5 min) and incubated in blocking buffer (BB, PBS containing 0.4% Fish Skin Gelatin and 0.2% Triton X) for 45 min to block non-specific antibody binding sites. Sections were then incubated with primary antibody diluted in BB for 1h at room temperature and overnight at 4°C. Mouse anti-dopamine β-hydroxylase (DBH, 1:1000, Merck) was used to visualize LC neurons and rabbit anti-red fluorescent protein to visualise the fRed tag (tRFP, 1:1000, Evrogen). For visualisation of GRAB_NE2m_ expression in the hippocampus, chicken anti-GFP (1:2000, Abcam) was used. A negative control containing BB without antibodies was used. The following day, sections were rinsed in blocking buffer (2x 10 min) and incubated with secondary antibodies: Alexa Fluor 488 goat anti-mouse (1:1000, Abcam), Alexa Fluor 594 goat anti-rabbit (1:1000, Abcam) and Alexa Fluor 488 goat anti-chicken (1:1000, Abcam) diluted in BB for 1h at room temperature in darkness. After rinsing in PBS (2x 5min), sections were mounted on glass slides and cover slipped with Vectashield H1000 mounting medium (Vector Laboratories, USA) to prevent photo bleaching.

## Acknowledgements

The research performed was supported by research grants from the Flemish Research Foundation (11M6422N) and the Queen Elisabeth Medical Foundation.

## Notes

### Competing Interest Statement

The authors have declared no competing interest.

### Summary of Updates

Corrected a couple of typos.

