## Supplementary Figures for "Modulation of locus coeruleus neurons and strong release of noradrenaline during acute hippocampal seizures in rats"

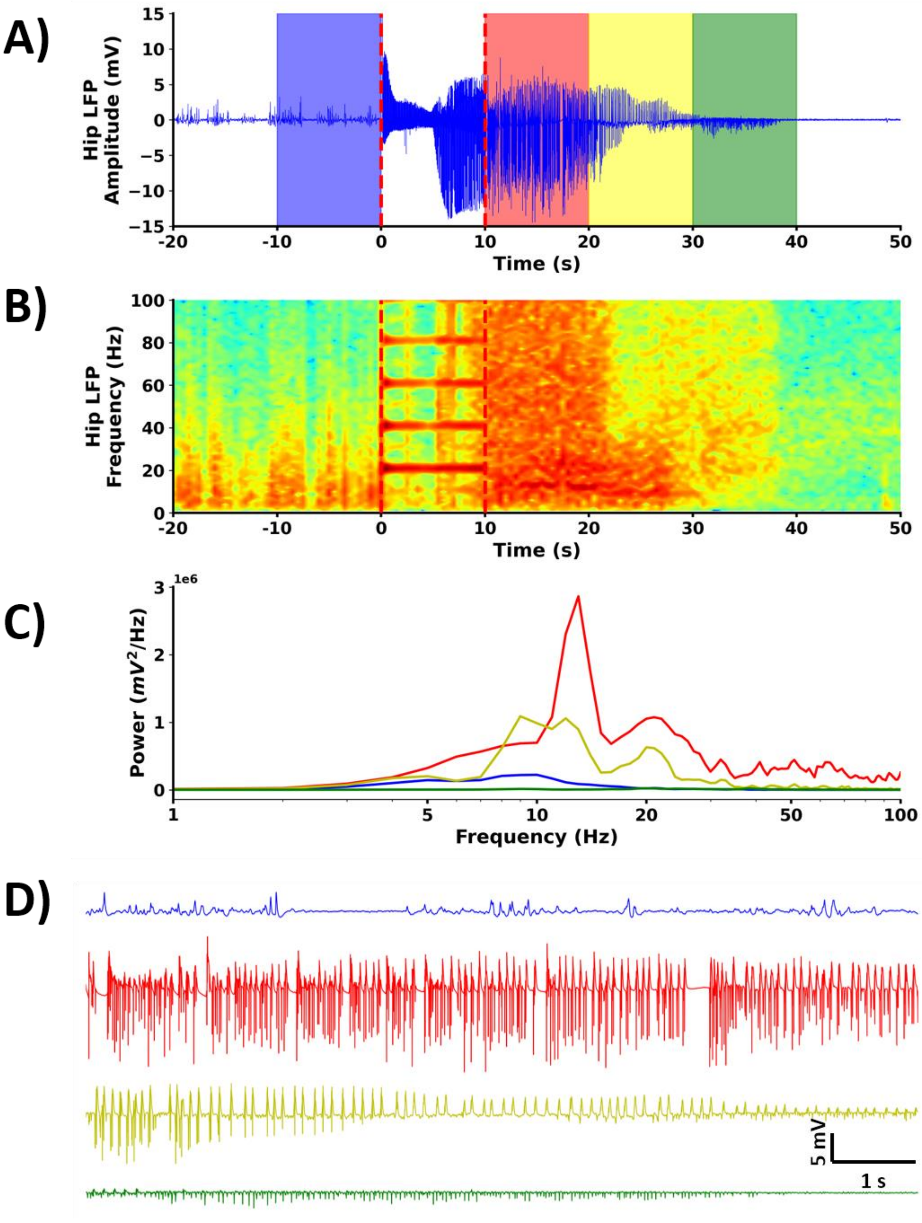


**Supplementary Figure 1:** Example of a hippocampal seizure evoked by perforant path stimulation. A) The Hippocampal local field potential (LFP) trace in mV over time, the red dotted lines indicate the 10s perforant path stimulation, B) the corresponding spectrogram, C) power spectra of 10 second epochs marked with shading in A), and D) a zoomed view on the 10 second traces with colors referring to the color of the shaded areas indicated in A).

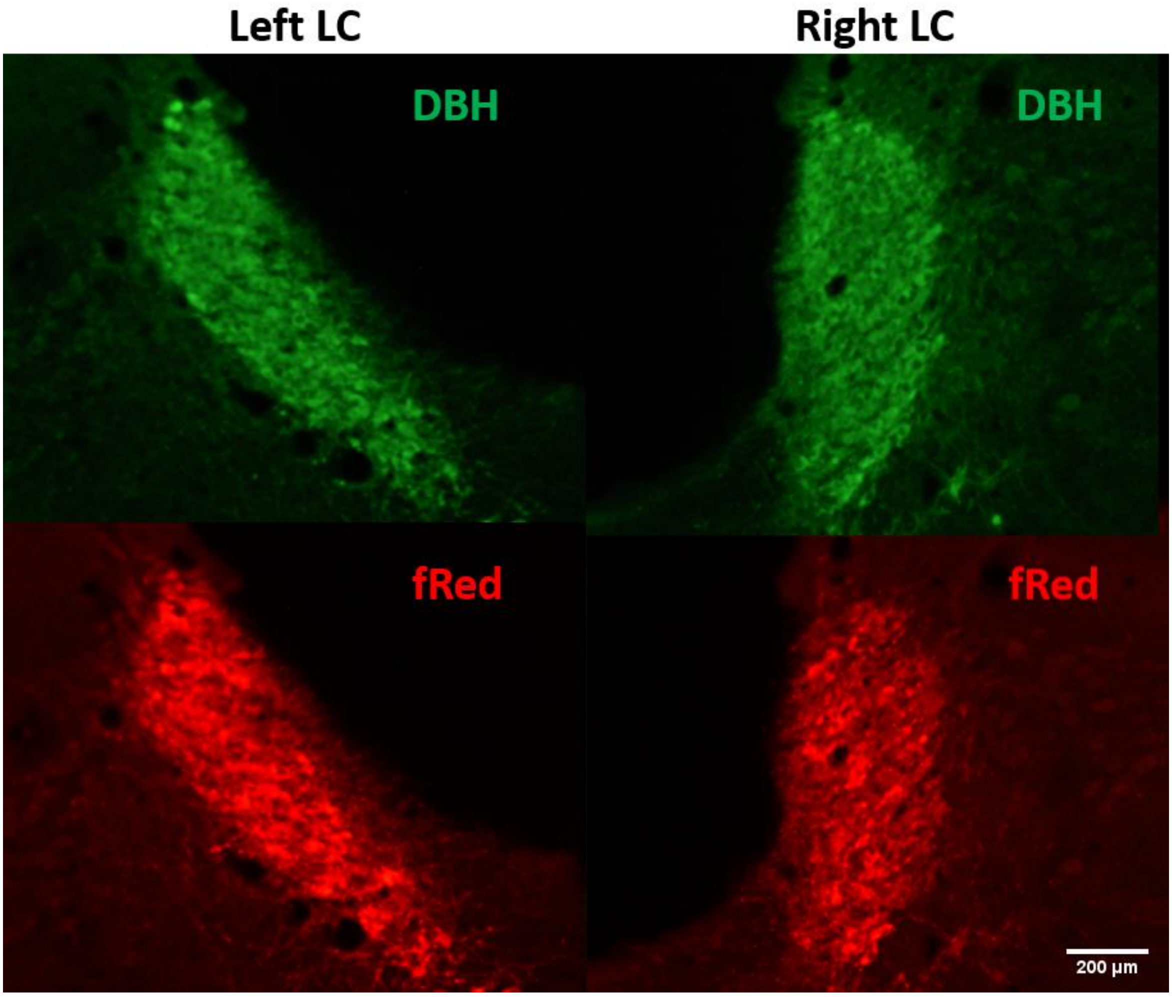


**Supplementary Figure 2:** Example of bilateral expression of the inhibitory opsin GtACR2 in the LC. Specific expression is indicated by a complete overlap between the $DBH^+$ neurons (primary antibody mouse anti-DBH, AF 488, green) and $fRed^+$ neurons (primary antibody rabbit anti-tRFP, AF 594, red), in both the left and right LC.

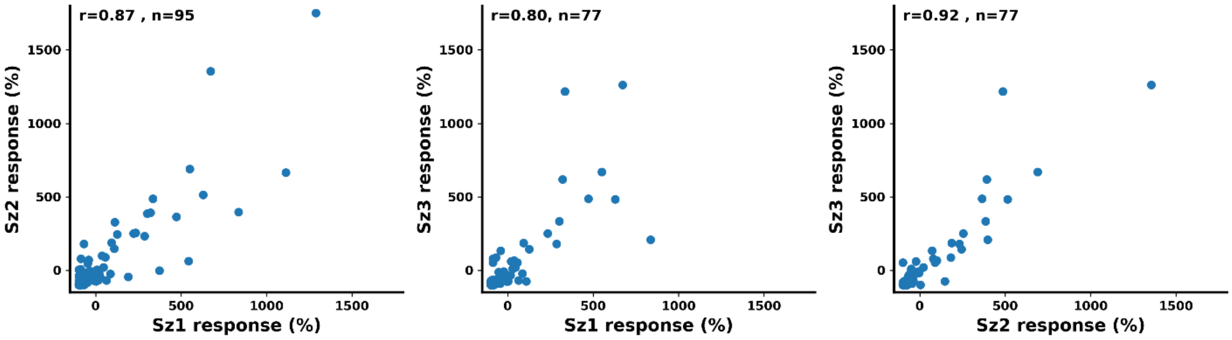


**Supplementary Figure 3:** Correlation of locus coeruleus (LC) neuronal responses to consecutively evoked hippocampal seizures (sz). The r-values provided denote the Spearman correlation coefficients and n represents the number of isolated LC neurons.

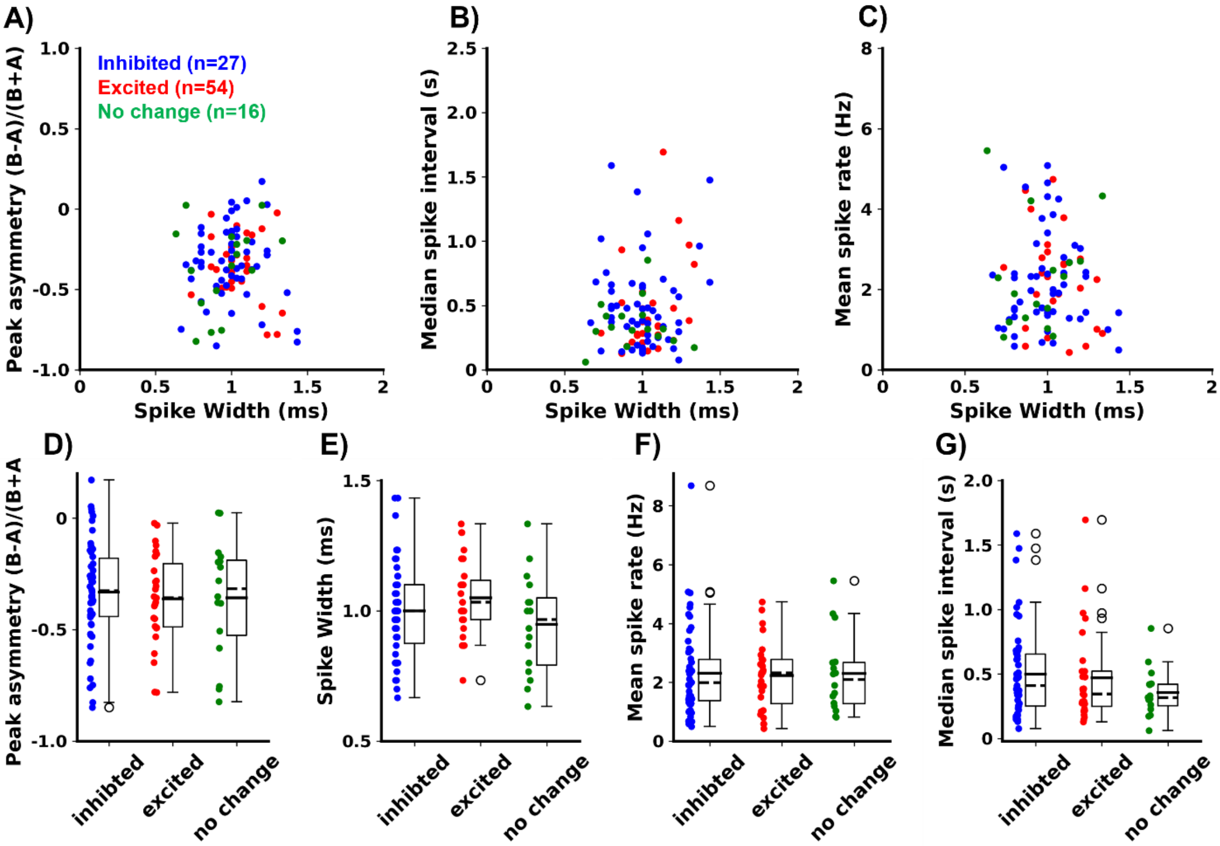


**Supplementary Figure 4:** Characteristics of spikes recorded from locus coeruleus (LC) neurons stratified for their response to evoked hippocampal seizures. A), B) and C) show no tendencies for clustering in spike characteristics, when assessing spike width, asymmetry of peaks in the spike waveforms, median spike interval or mean spike rate. D), E), F) and G) further show no differences in these variables when grouping as a function of how the neurons respond to seizures.

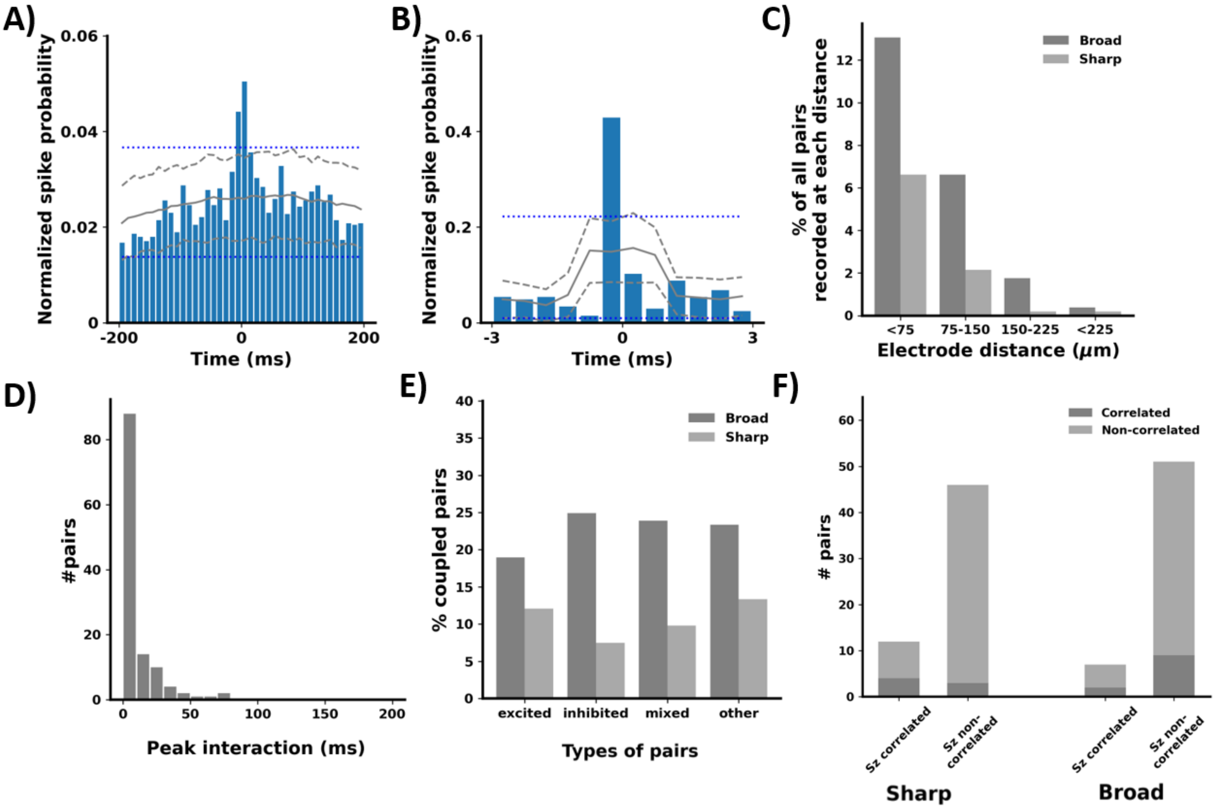


**Supplementary Figure 5:** Cross-correlation between pairs of locus coeruleus neurons in the interictal periods (n=513 pairs). As reported previously (Totah et al., 2018b), two types of cross-correlation were assessed: A) cross-correlation over a window of ±200 ms, called broad coupling and B) over a window of ±3 ms, called sharp coupling. C) The frequency of both broad and sharp coupling was related to the distance between electrode pairs of the electrode contacts where the spike amplitude was highest and was comparable to previous reports. D) For broad coupling, the peak of the interaction was most commonly observed around 0, as in the example in A). E) Although the previously reported coupling was observed, interictal coupling was observed at similar rates between pairs of neurons excited during seizures, inhibited neuron pairs and excited/inhibited neuron pairs. F) Finally, assessment of coupling of LC neurons during seizures showed that although not all neuron pairs with interictal coupling showed coupling during seizures, preictal sharp coupling was nevertheless observed to predict coupling during seizures.

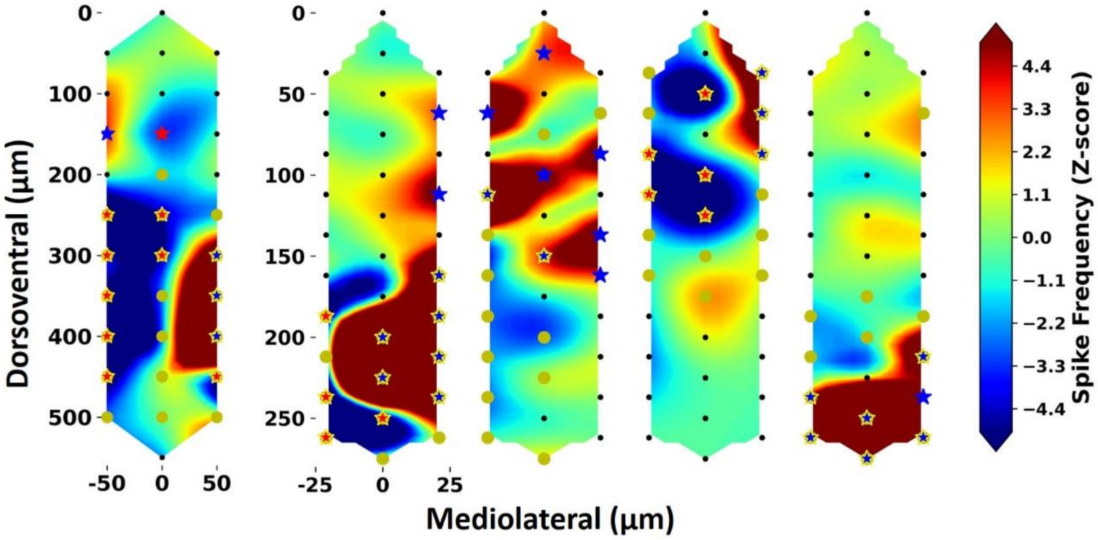


**Supplementary Figure 6:** Examples from different animals and different recordings of tendencies for anatomical separation of areas being inhibited vs. excited during hippocampal evoked seizures. Significant responses at an individual channel level are indicated with yellow dots. Red stars indicated significant inhibition and blue starts depict significant excitation. Channels recording LC neuronal activity, based on a significant burst inhibition in response to a noxious foot pinch, are noted with a yellow edge around the stars.

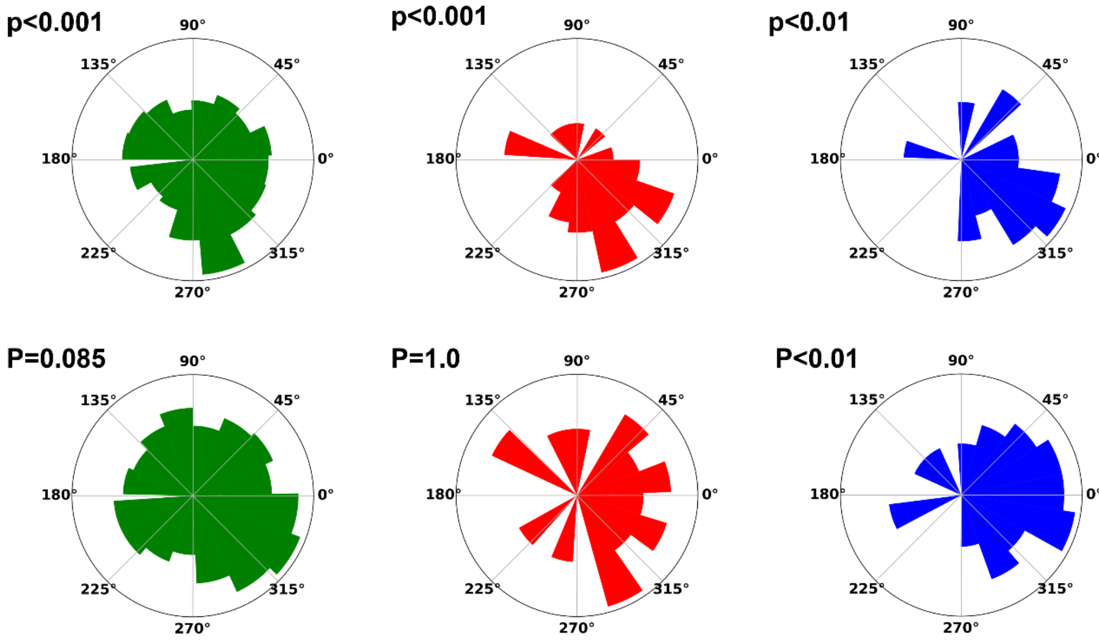


**Supplementary Figure 7:** Phase preference of three locus coeruleus (LC) neurons to the filtered (8-16 Hz) hippocampal local field potential, over two consecutive hippocampal seizures. On neuron (blue), retains its coupling, while the coupling of two other neurons (green and red) showed less consistency.

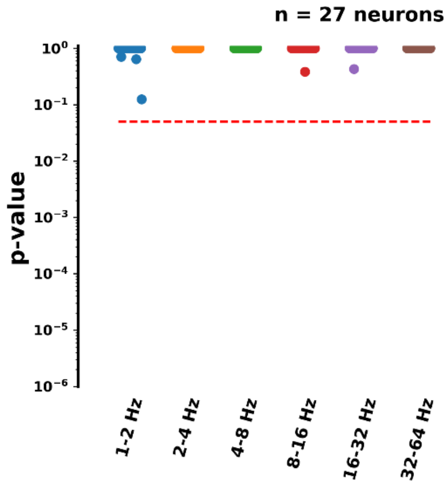


**Supplementary Figure 8:** Coupling of LC neurons to the phase of the bandpass filtered hippocampal local field potential (LFP) in the preictal period (60 seconds prior to seizure induction). None of the neurons showed any coupling to the hippocampal LFP.

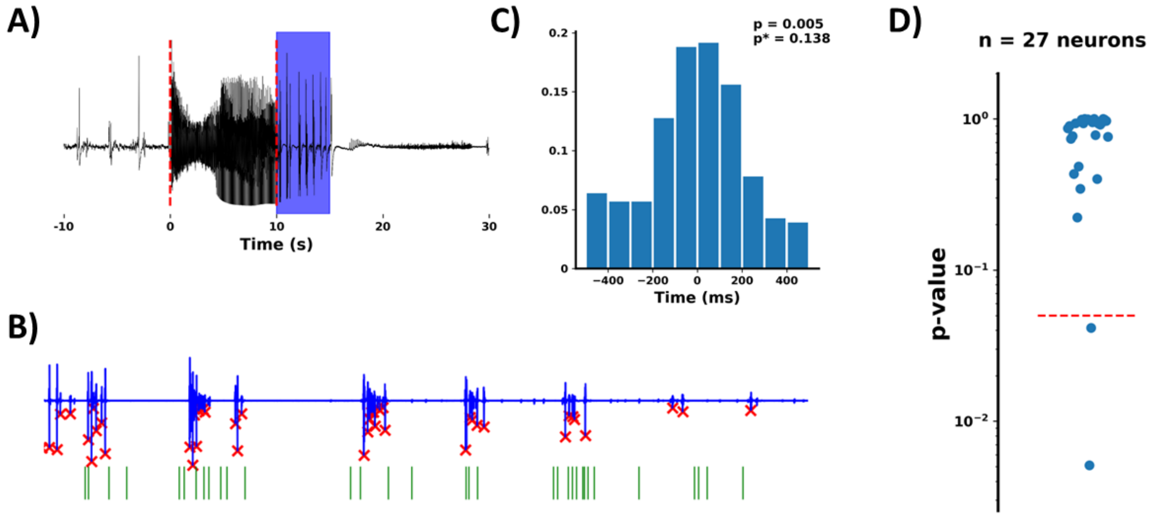


**Supplementary Figure 9:** Analysis of coupling between hippocampal seizures associated population spikes and locus coeruleus (LC) unit activity. A) Example of hippocampal seizure LFP, B) an excerpt of the hippocampal LFP marked with blue shading in A) and associated timing of LC unit showing the statistically strongest coupling to the population spike peaks (red crosses). C) Histogram of LC unit spike timings of neuron showing the strongest coupling, with an increased tendency for spiking around the negative peak of the population spike. This pattern reached statistical significance, which however did not remain following significance correction for multiple testing (p*). D) Distribution of uncorrected p-value outcomes of statistical assessment of coupling between hippocampal LFP population spikes in relation to seizures and LC spike timings of all 27 excited LC neurons.
